# Implementation-independent representation for deep convolutional neural networks and humans in processing faces

**DOI:** 10.1101/2020.06.26.171298

**Authors:** Yiying Song, Yukun Qu, Shan Xu, Jia Liu

## Abstract

Deep convolutional neural networks (DCNN) nowadays can match and even outperform human performance in challenging complex tasks. However, it remains unknown whether DCNNs achieve human-like performance through human-like processes; that is, do DCNNs use similar internal representations to achieve the task as humans? Here we applied a reverse-correlation method to reconstruct the internal representations when DCNNs and human observers classified genders of faces. We found that human observers and a DCNN pre-trained for face identification, VGG-Face, showed high similarity between their “classification images” in gender classification, suggesting similar critical information utilized in this task. Further analyses showed that the similarity of the representations was mainly observed at low spatial frequencies, which are critical for gender classification in human studies. Importantly, the prior task experience, which the VGG-Face was pre-trained for processing faces at the subordinate level (i.e., identification) as humans do, seemed necessary for such representational similarity, because AlexNet, a DCNN pre-trained to process objects at the basic level (i.e., categorization), succeeded in gender classification but relied on a completely different representation. In sum, although DCNNs and humans rely on different sets of hardware to process faces, they can use a similar representation, possibly from similar prior task experiences, to achieve the same computation goal. Therefore, our study provides the first empirical evidence supporting the hypothesis of implementation-independent representation.

## Introduction

In recent years, deep convolutional neural networks (DCNN) have made dramatic progresses to achieve human-level performances in a variety of challenging complex tasks, especially visual tasks. For example, DCNNs trained to classify over a million natural images can match human performance on object categorization tasks^1–3^, and DCNNs trained with large-scale face datasets can approach human-level performance in face recognition^4–7^. However, these highly complex networks have remained largely opaque, whose internal operations are poorly understood. Specifically, it remains unknown whether DCNNs achieve human-like performance through human-like processes. That is, do DCNNs use similar computations and inner representations to perform tasks as humans do?

To address this question, here we applied a reverse correlation approach^8–11^, which uses outputs (e.g., behavior performance) to infer internal representations that transform inputs (e.g., stimuli) to outputs for DCNNs and human observers. This data-driven method allows an unbiased estimate of what is in observers’ “mind” when performing a task, rather than manipulates specific features that researchers *a priori* hypothesize to be critical for the task. Here we investigated whether the DCNNs and humans utilized similar representations to perform the task of face gender classification.

Specifically, a gender-neutral template face midway between the average male and the average female faces was superimposed with random noises, which rendered the template face more male-like in some trials or more female-like in other trials. The noisy faces were then submitted to human observers and the VGG-Face, a typical DCNN pre-trained for face identification^4^. Based on the output of an observer that a noisy face was classified as a male but not as a female, for example, we reasoned that the noise superimposed on the template face contained features matching the observer’s internal male prototype. Therefore, the difference between noise patterns of trials classified as male and those as female provided an explicit and unbiased estimate of the representation used by the observer for gender classification. Finally, we directly compared the similarity of the inner representations of human observers and the VGG-Face obtained from identical stimuli and procedures, and examined the hypothesis that different intelligent information-processing systems may use similar representations to achieve the same computation goal^12^.

## Results

We used the reverse correlation approach to reconstruct the inner representations used by the DCNN and human observers for gender classification. Specifically, both the DCNN and human observers were asked to classify noisy faces from a gender-neutral template face embedded with random sinusoid noises as male or female (Figure 1a).

**Figure 1.**
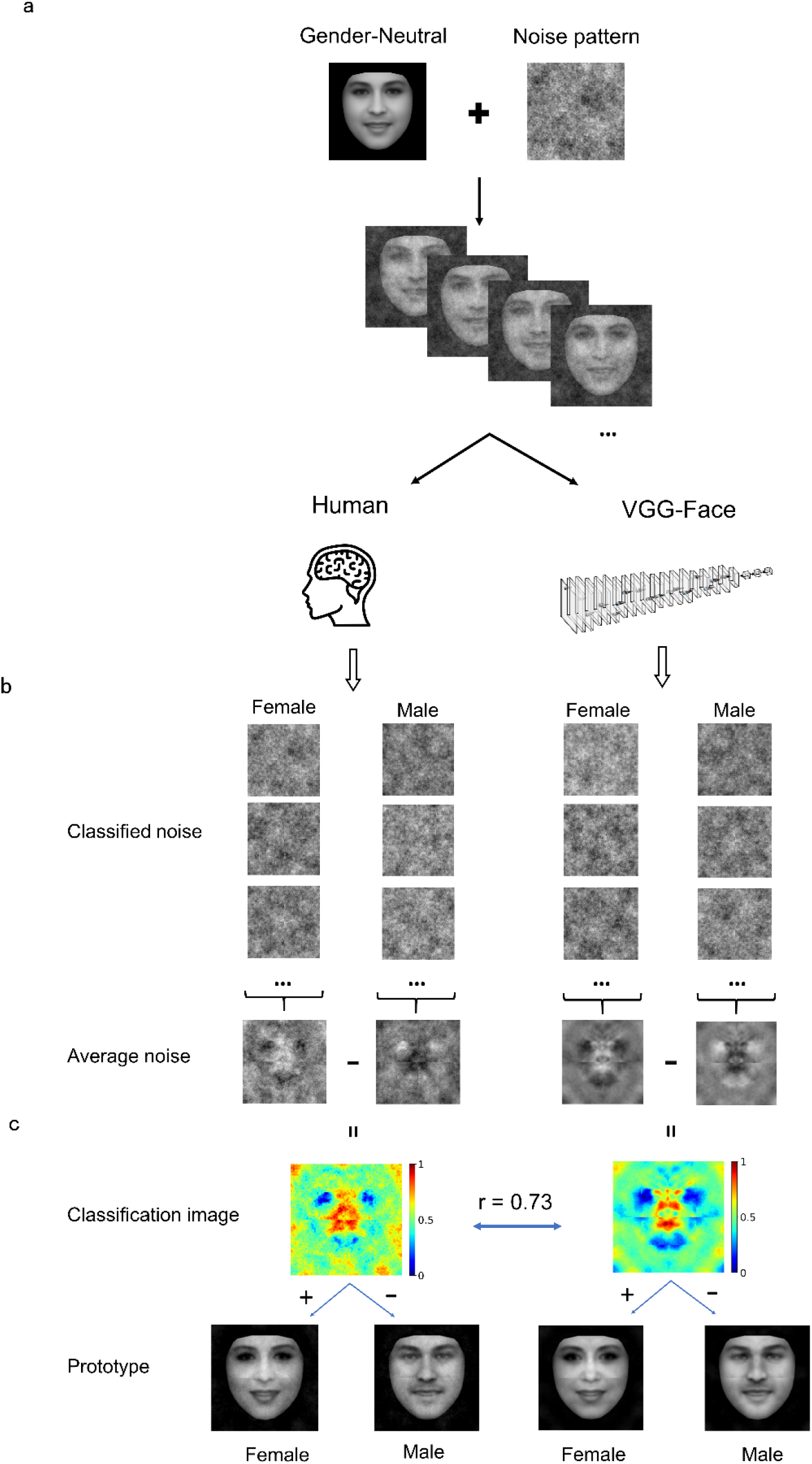
(a) Experiment procedure. A gender-neutral template face was superimposed with noises to create a set of gender-ambiguous faces, which were submitted to the VGG-Face and human observers for gender classification. (b) Exemplars of noises extracted from noisy faces classified as either female or male, respectively. The noises were then averaged to reconstruct images that contained the critical information for classifying the noisy faces as male or as female. (c) Classification images (CI) was the contrast of the average noise of female by that of male. For visualization, values in each CI were normalized separately to the range from 0 to 1, denoted by colors. By adding or subtracting the rescaled CI to or from the gender-neutral template face, female or male prototype of human observers (Left) and the VGG-Face (Right) were created. The faces shown in the figure were synthetic ones created by a morph algorithm and thus had no relation to real human.

For the DCNN, we first trained the VGG-Face to classify gender using transfer learning with 21,458 face images of 52 identities (35 males) from the VGG-Face2 dataset (see Method), and the test accuracy of gender classification of the new network achieved 100%. The gender-neutral template face was roughly equally classified as male and female by the VGG-Face (female: 54%). The noise patterns were constructed from 4,092 sinusoids at five spatial scales, six orientations, and two phases. We presented the template face embedded in 20,000 noise patterns to the VGG-Face, of which 11,736 (58.7%) images were classified as male and 8,264 (41.3%) images as female. The noise patterns from trials classified as male or female were averaged separately (Figure 1b), and the difference between the two average noise patterns yielded a “classification image” (CI) that makes explicit the information used by the VGG-Face for gender classification (Figure 1c). A visual inspection of the CI showed that regions around the eyes, nose, and mouth were of high contrast in the CI, indicating the critical regions employed by the VGG-Face to classify male from female faces.

Then, we reconstructed the representation used by human observers in a similar way. In our study, 16 human observers performed the gender classification task, each presented with 1000 noisy faces. Altogether, 16,000 images were presented to the human observers, of which 7,969 (49.8%) images were classified as males and 8,031(50.2%) images as females. Similarly, the CI for human observers was obtained (Figure 1c). Visual inspections of the CIs for the VGG-Face and human observers revealed good agreement between them, and Pearson’s correlation between the two CIs was high (*r* = 0.73). This result suggested that the VGG-Face and human observers utilized similar information to classifying gender.

Further, we reconstructed inner male and female prototypes by adding or subtracting the rescaled CI to or from the template face for the VGG-Face and humans respectively (Figure 1c). As expected, the male and female prototype faces are perceptually male-like and female-like, and highly similar between the VGG-Face and human observers.

Having found that the VGG-Face and human observers utilized similar information for gender classification, next we asked whether the VGG-Face and human observers employed similar information in all spatial frequencies. In our study, the noise patterns were constructed from sinusoid components of five scales of spatial frequencies (2, 4, 8, 16, and 32 cycles/image), which enabled us to reconstruct the CIs for each scale separately (Figure 2) and examined the similarity at each scale. We found that the similarity was the highest at low spatial frequencies (*r* = 0.87 and 0.76 at 2 and 4 cycles/images), and then decreased sharply at high spatial frequencies (*r* = 0.25, 0.19, 0.11 at 8, 16, and 32 cycles/image). Consequently, male and female prototypes reconstructed with the noise patterns at low spatial frequencies (2 and 4 cycles/image) were more similar between human observers and the VGG-Face than those at high spatial frequencies (8, 16, and 32 cycles/images) (Supplementary Analysis 1). Therefore, the shared representation for gender classification was mainly based on information at low spatial frequencies, consistent with previous findings that face gender processing relies heavily on low spatial frequencies^9, 13–16^.

**Figure 2.**
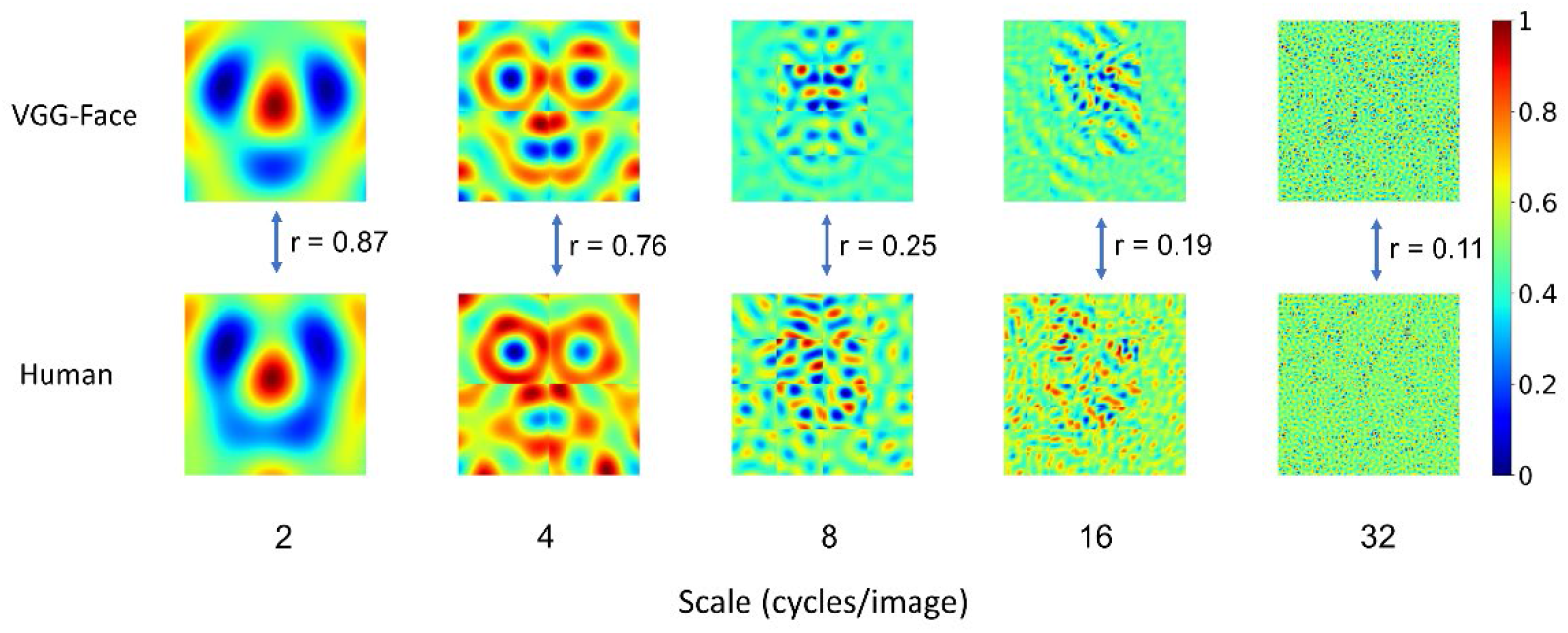
Correspondence in representation at different scales of spatial frequencies. For visualization, values in each CI were normalized separately to the range from 0 to 1, denoted by colors. Note that the correspondence was the highest at the low-spatial frequencies, and then decreased sharply at the high-spatial frequencies. Scale number denotes cycles per image.

To further quantify the contribution of different spatial frequencies for gender classification, we calculated the contribution of each of the 4,092 parameters from all five spatial frequencies. For each parameter, we performed an independent sample t-test (two-sided) between the parameter values from the male trials and those from the female trials, and calculated the absolute value of Cohen’s d as an index of the contribution of each parameter to gender classification. One hundred and four parameters in the VGG-Face and 12 in human observers contributed significantly for the classification (Bonferroni corrected for multiple comparisons, Figure 3a and b). Of the 12 parameters in human observers, 9 were at the scales of 2 and 4 cycles/images. Similarly, most of the 104 parameters in the VGG-Face were also at low-frequency scales (7 at 2 cycles/images, 33 at 4 cycles/images, and 30 at 8 cycles/images), and the percentage of the significant parameters at low frequencies (58% and 69% at 2 and 4 cycles/images) were much higher than those at high frequencies (4% and 0% at 16 and 32 cycles/images). That is, both the VGG-Face and human observers mainly relied on information at low spatial frequencies for gender classification.

**Figure 3.**
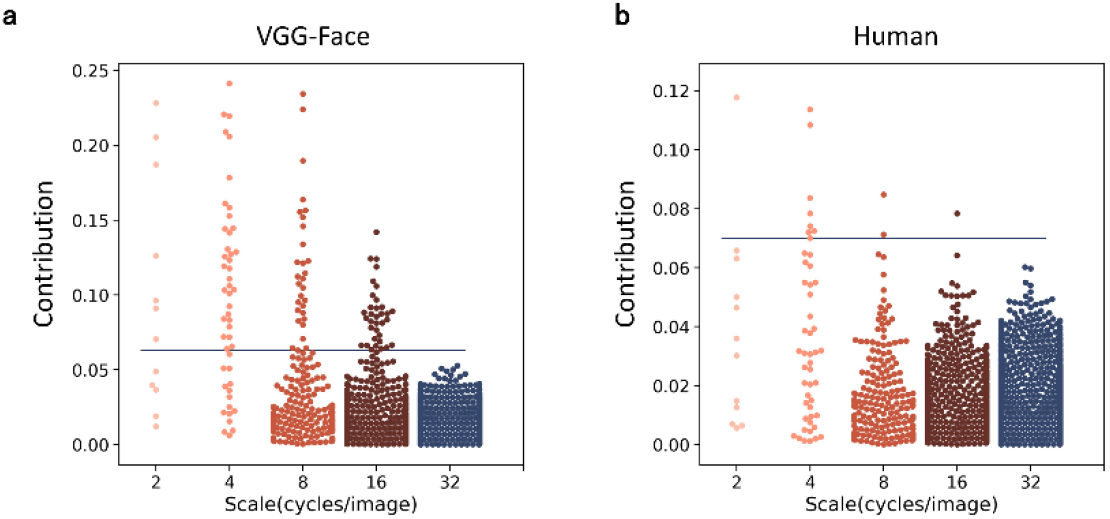
Manhattan plot of the contribution (the absolute value of Cohen’s d) of the parameters used to construct noises in the VGG-Face (a) and in human observers (b). Each dot denotes a parameter, and the horizontal blue line indicates the significance level after Bonferroni correction.

Another way is to select parameters that made the most contributions indexed by the absolute values of Cohen’s d. We found that the 1,885 most contributing parameters of all 4,092 parameters already made up to 80% of the total contribution for the VGG-Face; importantly, these parameters also made up 48% of the contribution for human observers. Then, we examined the similarity of parameters’ contribution by calculating the Spearman’s correlation between Cohen’s d of the VGG-Face and human observers for the highly-contributing parameters at each scale of spatial frequencies. We found that the correlation was high at low spatial frequencies (*r* = 0.79 and 0.74 at 2 and 4 cycles/images), and then declined sharply at high spatial frequencies (*r* = 0.21, 0.27, and 0.17 at 8, 16, and 32 cycles/images). In contrast, there were more parameters at high than low spatial frequencies that contributed differently between the VGG-Face and human observers (Supplementary Analysis 2). Taken together, at low spatial frequencies, not only were the representations more similar, but also the parameters underlying the representation contributed more significantly to the task.

Where did the representational similarity come from? One possibility is that information at low spatial frequencies is critical for face processing, and therefore both DCNN and human observers were forced to exact information at low spatial frequencies to successfully perform the task. An alternative hypothesis is that the VGG-Face and human observers share similar prior experiences of processing face at the subordinate level where faces are identified into different individuals. To test these two hypotheses, we examined another typical DCNN, the AlexNet, that also has abundant exposure to face images but is pre-trained to classify objects into 1000 basic categories. We trained the AlexNet to perform the gender classification task with the same transfer learning procedure as that for the VGG-Face. The testing accuracy of gender classification of the AlexNet reached 93%, indicating that it was able to perform the task. However, the CI obtained from the AlexNet (Figure 4a) was in sharp contrast to the CIs of human observers (Figure 1c) as a whole (*r* = −0.04) and at different scales (*r* = −0.28, 0.03, 0.25, 0.10 and 0.03 at the scales of 2, 4, 8, 16 and 32). We also reconstructed the female and male prototype faces of AlexNet (Figure 4a), and they appeared quite distinct from those of human observers and the VGG-Face (Figure 1c). This finding was unlikely due to the differences in architecture between the VGG-Face and the AlexNet, because the VGG-16, which has the same architecture as the VGG-Face but is pre-trained for object categorization as the AlexNet, also showed a CI largely different from human observers (Supplemental Analysis 3). Therefore, although the AlexNet succeeded in performing the gender classification task, it relied on a set of information completely different from human observers to achieve the goal. Therefore, mere exposure to face stimuli is not sufficient for the DCNNs to construct similar representations for gender classification as human observers; instead, the prior experience of face identification was required.

**Figure 4.**
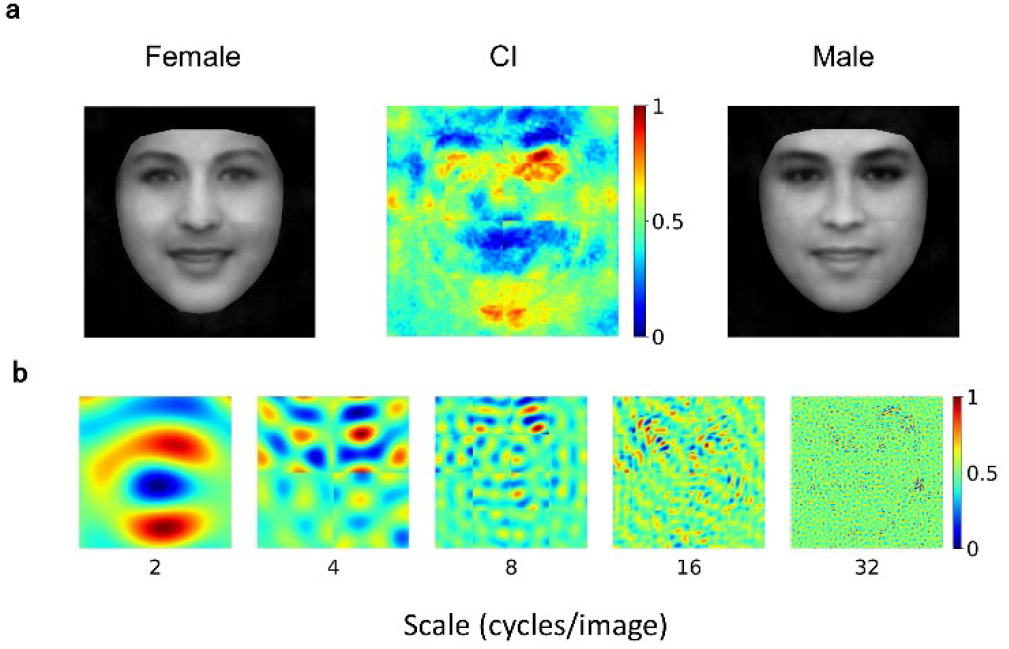
(a) AlextNet’s CI for gender classification. For visualization, values in each CI were normalized separately to the range from 0 to 1, denoted by colors. Note that the female prototype (left) and the male prototype (right) were not perceptually female-like and male-like respectively. (b) Normalized CI at different scales of spatial frequencies. Note that they were significantly different from those of human observers. The faces shown in the figure were synthetic ones created by a morph algorithm and thus had no relation to real human.

## Discussion

Marr (1982) has proposed a three-level framework to understand an intelligent information-processing system^12^. At the top is the computational level that defines the goal of the system, and in our study, the computation goal is face gender classification; at the bottom is the implementation level that is the physical substrate of the system, which are the DCNNs and human brain in our study. Most critically, in the middle is the representational and algorithmic level that establishes approaches through which the implementation achieves the computation goal. Despite dramatic differences in the physical implementations between the artificial and biological intelligent systems, similar representations may be used by different systems to achieve the same computation goal. Our study provides one of the first direct evidence to support this hypothesis by showing that the DCNNs and humans used similar representations to achieve the goal of face gender classification, which were revealed by highly similar CIs between the VGG-Face and humans.

The shared representation, on one hand, may come from the critical stimulus information needed to achieve the computation goal. Previous human studies on gender classification suggest that the critical information humans used to solve the task is embedded mainly in low spatial frequencies^9, 13–16^. Here we found that the VGG-Face also relied heavily on low spatial frequencies of faces for gender classification. Further, it was the information only in this band that showed similarity to that of humans, but not in high spatial frequencies. In other words, one reason that the VGG-Face and humans established similar representations based on low spatial frequencies might be that this stimulus information is critical for the task of face gender classification.

On the other hand, the prior task experience before the gender classification task may also play a deterministic role for DCNNs to use a similar approach to achieve the goal as humans. Previous studies have shown that humans usually process faces at the subordinate level, that is, to recognize faces as individuals. Similar to humans, the VGG-Face is also pre-trained to recognize faces at the individual level, that is, to classify face images into different identities (e.g., John’s face). Therefore, the similar task experience in the past likely led the similar approaches in achieving the new goal of gender classification.

In contrast, the AlexNet is pre-trained to recognize objects at the basic level, that is, to classify objects into categories (e.g., dogs) but not individuals (John’s dog). Therefore, although the AlexNet experiences abundant exposure to face images during the pre-training, it processes faces as objects, different from humans and the VGG-Face. Previous studies on humans have shown that object recognition does not selectively rely on low- to middle- spatial frequencies as face recognition does^17–20^. Thus, it is not surprising that although the AlexNet achieved a comparable performance in face gender classification as humans, an approach significantly different from that of humans was adopted. Taken together, the similarity in representation between DCNNs and humans was not guaranteed by the common computational goal or by the passive experiences with stimuli; instead, it was constrained by the combination of experiences on the pre-training task in the past and critical stimulus information needed in performing the task in the present.

In history, two main approaches have been proposed to achieve and even excel human vision in artificial intelligence^21^. The neuroscience approach adheres to biological fidelity at the implementation level, which simulates neural circuits of brains, whereas the cognitive approach emphasizes on cognitive fidelity, which focuses on goal-directed algorithms and disregards implementation. Our study suggests an intermediate approach lying in between these two. By simulating human intelligence at the representation level in Marr’s framework, this approach provides an abstract description of how a system extracts critical features to construct representation for a specific task. Because the representation is relatively independent of implementation, the knowledge acquired in biological systems can be easily adopted by artificial systems with completely different substrates. Therefore, the simulation of representation may shed light on building new AI systems in a feasible way.

## Methods

### Transfer learning

We used the pre-trained VGG-Face network^4^ that consists of 13 convolutional layers and 3 fully connected (FC) layers. Each convolutional layer and FC layer were followed by one or more non-linearities such as ReLU and max pooling. The VGG-Face network was pre-trained for face identification with the VGG-Face dataset containing over two million face images of 2,662 identities.

In our study, we trained the VGG-Face for face-gender classification using transfer learning. The final FC layer of the VGG-Face has 2,662 units, each for one identity. We replaced this layer with a two-unit FC layer for the binary gender classification. All weights of the network were frozen except the weights between the penultimate FC layer and the new final FC layer. The training sample contains 21,458 face images (male: 14,586) of 52 identities (male: 35) randomly selected from the VGG-Face2 dataset. The testing sample contains other 666 face images (male: 429) from the same 52 identities. All face images were resized to 224×224 pixels to match the model input size. We used in-house python package DCNNbrain to train the network. The loss function was cross-entropy, and the optimizer was Adam. The learning rate was 0.03, and the network was trained for 25 epochs. After training, the accuracy of gender classification on the testing sample reached 100%.

The same training procedures were applied to AlexNet pre-trained for object categorization^2^. The model consists of five convolutional layers and three FC layers. The AlexNet was pre-trained on ImageNet to classify 1.2 million images into 1000 object categories. In our study, after transfer learning, the accuracy on the testing sample reached 93% for gender classification.

### Reverse correlation Approach

After the transfer learning on gender classification, we made the representation explicit with the reverse correlation approach on noisy faces. All stimuli consisted of a gender-neutral template face superimposed with sinusoid noise patterns. The template was a morphed face between a female average face and a male average face (Figure 1a). The female and male average faces were computed as a mathematical average of all female and all male faces of the training sample after they were aligned and wrapped into the same space with 68 landmarks using an open-access toolbox face_morpher (https://github.com/alyssaq/face_morpher). The average faces were 255-bit grayscale and 512 × 512 pixel images. We further created 500 morph faces that gradually changed from the female average face to the male average face using face_morpher. Then we presented 500 morphed faces evenly distributed between the female and the male average faces to the VGG-Face to find the face most equally classified as male and female in gender classification. The 250th morphed face, which was classified as female with a probability of 54% was chosen as the gender-neutral template face in our study.

A random noise pattern was generated for each trial. Each noise pattern was composed of sinusoid patch layers of five different scales of spatial frequencies (2, 4, 8, 16, 32 cycles/image), with each patch layer made up of 1, 4, 16, 64 and 256 sinusoid patches respectively^9^. For each sinusoid patch, sinusoids of six orientations (0, 30, 60, 90, 120, and 150 degrees) and two phases (0 and pi/2) were summed. The amplitude of each sinusoid came from a random sampling of a uniform distribution of values from −1 to 1. Therefore, each noise pattern was determined by 4,092 random amplitude parameters (12, 48, 192, 768, and 3072 parameters for 2, 4, 8, 16, and 32 cycles/image). We use the R package rcicr to generate the sinusoid noises^22^. We created 20,000 noise patterns for the DCNNs and 1,000 noise patterns for each human observer. Each noise pattern was then superimposed on the template face to create a different noisy face.

We resized the noisy face images to 224 × 224 pixels and submitted them to the VGG-Face and AlexNet, and obtained their classification prediction for each image. For VGG-Face, a noisy face was classified as male when the activation of the male unit was higher than the female unit. Note that the AlexNet showed a bias toward male faces when classifying the noisy faces; therefore we modified the classification criterion for the AlexNet. That is, for AlexNet, a noisy face would be classified as male when the activation of the male unit to the to-be-classified face was higher than its average activation to all noisy faces. Note that the choice of criterion would not affect the results pattern of the VGG-Face and hence the dissociation between AlexNet and VGG-Face, because the two criteria lead to literally identical CIs for VGG-Face (*r* = 0.99).

To generate corresponding female or male prototype faces, each CI was separately rescaled to have the same maximum pixel value and then added or subtracted from the template face.

### Participants

Sixteen college students (12 females, age 19 – 33 years, mean age 22 years) from Beijing Normal University, Beijing, China, participated in the gender classification task. All participants were right-handed and had normal or corrected-to-normal vision. The experiment protocol was approved by the Institutional Review Board of the Faculty of Psychology, Beijing Normal University. Written informed consent was obtained from all participants before the experiment.

### Experimental procedures

Before the experiment, participants were told that they would perform a difficult gender classification task because the faces were superimposed with heavy noises. The stimuli were 255-bit grayscale and 512 × 512 pixel images. PsychoPy^23^ was used to display the stimuli and record responses. The stimuli were presented on the screen of a Dell precision laptop at a distance of 70 cm. The stimuli subtended a visual angle of approximately 8.2 degree. In each trial, a noisy face image was presented in the center of the screen for 1s, and then the screen cleared until the participant made a response. The participants were instructed to provide one of four responses with a key press for each trial: probably female, possibly female, possibly male, or probably male. No feedback was provided. Each participant performed 1000 trials. The participants could rest every 100 trials. The total experiment duration was about one hour for each participant. In data analysis, the CI was calculated by subtracting the average noise patterns from all trials classified as male (probably male and possibly male) from those classified as female (probably female and possibly female).

## Supporting information

Supplemental results

## Data availability

The datasets generated during and/or analysed during the current study are available from the corresponding author on reasonable request.

## Code availability

The code used to generate results that are reported in the paper are available from the corresponding author on reasonable request.

## Notes

### Competing Interest Statement

The authors have declared no competing interest.

## References

1. Krizhevsky, A. One weird trick for parallelizing convolutional neural networks. arXiv preprint arXiv:1404.5997 (2014).

2. Krizhevsky, A., Sutskever, I. & Hinton, G.E. Imagenet classification with deep convolutional neural networks. in Advances in neural information processing systems 1097–1105 (2012).

3. Simonyan, K. & Zisserman, A. Very deep convolutional networks for large-scale image recognition. arXiv preprint arXiv:1409.1556 (2014).

4. Parkhi, O.M., Vedaldi, A. & Zisserman, A. Deep face recognition. (2015).

5. Taigman, Y., Yang, M., Ranzato, M.A. & Wolf, L. Deepface: Closing the gap to human-level performance in face verification. in Proceedings of the IEEE conference on computer vision and pattern recognition 1701–1708 (2014).

6. Schroff, F., Kalenichenko, D. & Philbin, J. Facenet: A unified embedding for face recognition and clustering. in Proceedings of the IEEE conference on computer vision and pattern recognition 815–823 (2015).

7. Ranjan, R., Sankaranarayanan, S., Castillo, C.D. & Chellappa, R. An all-in-one convolutional neural network for face analysis. in 2017 12th IEEE International Conference on Automatic Face & Gesture Recognition (FG 2017) 17–24 (IEEE, 2017).

8. Ahumada Jr, A. & Lovell, J. Stimulus features in signal detection. The Journal of the Acoustical Society of America 49, 1751–1756 (1971).

9. Mangini, M.C. & Biederman, I. Making the ineffable explicit: Estimating the information employed for face classifications. Cognitive Science 28, 209–226 (2004).

10. Gold, J.M., Murray, R.F., Bennett, P.J. & Sekuler, A.B. Deriving behavioural receptive fields for visually completed contours. Current Biology 10, 663–666 (2000).

11. Martin-Malivel, J., Mangini, M.C., Fagot, J. & Biederman, I. Do humans and baboons use the same information when categorizing human and baboon faces? Psychological Science 17, 599–607 (2006).

12. Marr, D. Vision: A computational investigation into the human representation and processing of visual information, henry holt and co. Inc., New York, NY 2 (1982).

13. Sergent, J. Microgenesis of face perception. in Aspects of face processing 17–33 (Springer, 1986).

14. Valentin, D., Abdi, H. & O’toole, A.J. Categorization and identification of human face images by neural networks: A review of the linear autoassociative and principal component approaches. Journal of biological systems 2, 413–429 (1994).

15. Khalid, S., Finkbeiner, M., König, P. & Ansorge, U. Subcortical human face processing? Evidence from masked priming. Journal of Experimental Psychology: Human Perception and Performance 39, 989 (2013).

16. Goffaux, V., Jemel, B., Jacques, C., Rossion, B. & Schyns, P.G. ERP evidence for task modulations on face perceptual processing at different spatial scales. Cognitive Science 27, 313–325 (2003).

17. Goffaux, V., Gauthier, I. & Rossion, B. Spatial scale contribution to early visual differences between face and object processing. Cognitive Brain Research 16, 416–424 (2003).

18. Collin, C.A., Therrien, M.E., Campbell, K.B. & Hamm, J.P. Effects of band-pass spatial frequency filtering of face and object images on the amplitude of N170. Perception 41, 717–732 (2012).

19. Biederman, I. & Kalocsais, P. Neurocomputational bases of object and face recognition. Philosophical Transactions of the Royal Society of London. Series B: Biological Sciences 352, 1203–1219 (1997).

20. Collin, C.A. Spatial-frequency thresholds for object categorisation at basic and subordinate levels. Perception 35, 41–52 (2006).

21. Kriegeskorte, N. & Douglas, P.K. Cognitive computational neuroscience. Nature neuroscience 21, 1148–1160 (2018).

22. Dotsch, R. rcicr: Reverse-correlation image-classification toolbox (R package version 0.4. 0). (2017).

23. Peirce, J., et al. PsychoPy2: Experiments in behavior made easy. Behavior Research Methods 51, 195–203 (2019).

