## Supplemental results for "Implementation-independent representation for deep convolutional neural networks and humans in processing faces"

### Supplementary materials

#### Supplementary analysis 1

To investigate how different scales of spatial frequencies affected gender classification, we constructed male and female prototypes for each spatial frequency respectively. Visual inspection of Figure S1a revealed that the prototype images at 2 and 4 cycles/images were highly similar between the VGG-Face and human observers, while those at 8 and 16 cycles/images showed noticeable differences. To validate this intuition, we presented these prototype images to the VGG-Face, and examined whether they elicited similar activations in the final fully-connected layer. We found that for the scales at low spatial frequencies (2 and 4 cycles/images), the prototype images obtained from the VGG-Face and human observers elicited similar activation amplitudes, while at the scales of high spatial frequencies (8, 16, and 32 cycles/images), the prototype images obtained from the VGG-Face elicited much higher activation amplitudes than those obtained from humans (Figure S1b). Taken together, the VGG-Face and human observers shared similar inner representations at low spatial frequencies, and the similarity decreased along with the increase of spatial frequencies.

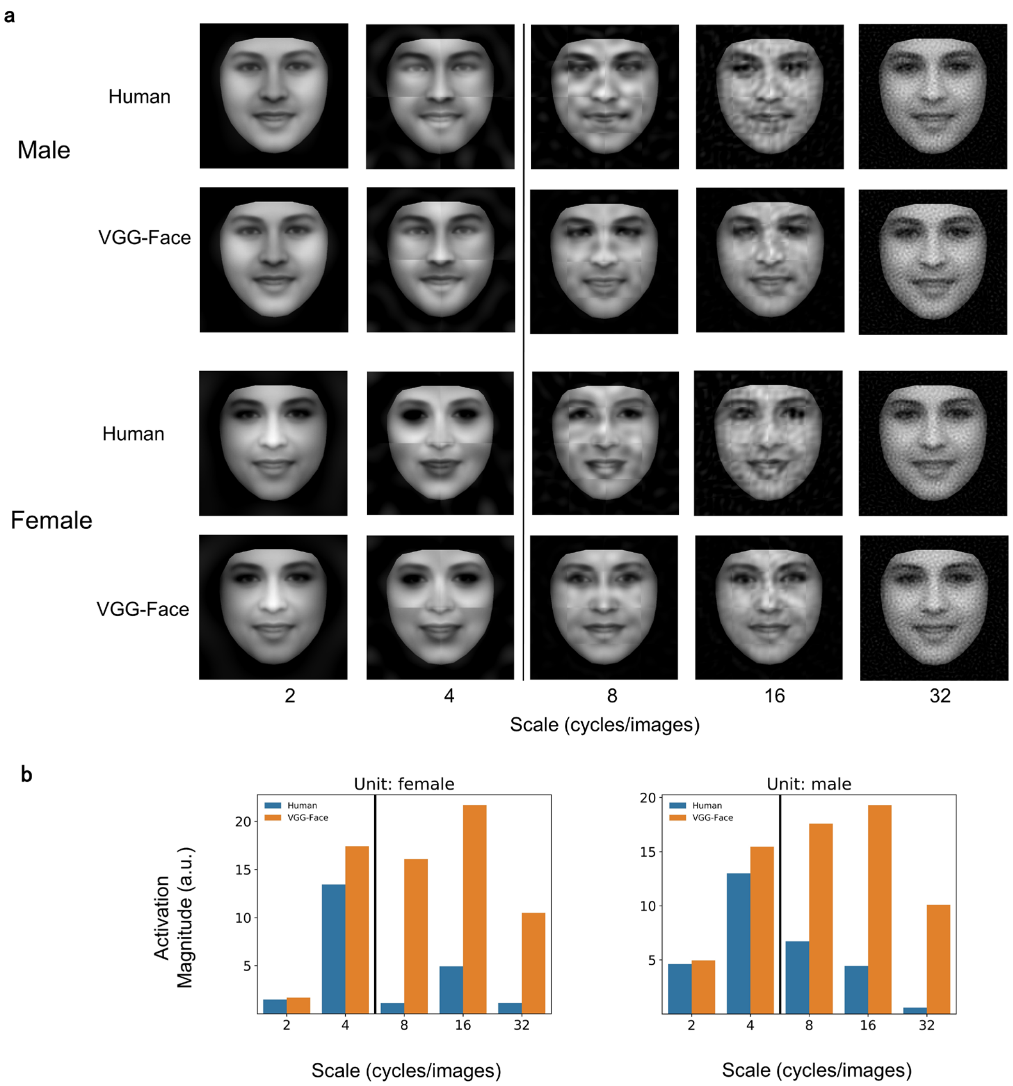


**Figure S1**. Male and female prototypes at different scales. (a) male and female prototypes of the VGG-Face and human observers at different scales of spatial frequencies. (b) the activation magnitude of the units at the output layer of the VGG-Face (Left: female; Right: male).

### Supplementary analysis 2

To examine the parameters that contributed differently between the VGG-Face and human observers, we first standardized the Cohen's d, the index for the size of contribution, to z-score for the VGG-Face and human observers respectively so that we can compare the contribution of the parameters across different scales. That is, the larger difference between the two z-scores of a parameter was, the more distinct contribution of the parameter between the VGG-Face and human observers. When the difference of a parameter deviated from the average contribution difference more than 1.96 standard deviations of the distribution of contribution differences, the difference was defined as significant. As shown in Figure S2a, the number of parameters that contributed significantly different between the VGG-Face and human observers differed. Further analyses revealed that the number of the parameters increased monotonically as a function of the scales of spatial frequencies, which were 3, 21, 40, 55, 64 at the scales of 2, 4, 8, 16, and 32 cycles/image respectively (Figure S2b). This result supplemented the finding that the VGG-Face and human observers made similar use of low spatial frequencies by showing that the difference in representation mainly came from face information at high spatial frequencies. The faces shown in the figure were synthetic ones created by a morph algorithm and thus had no relation to real human.

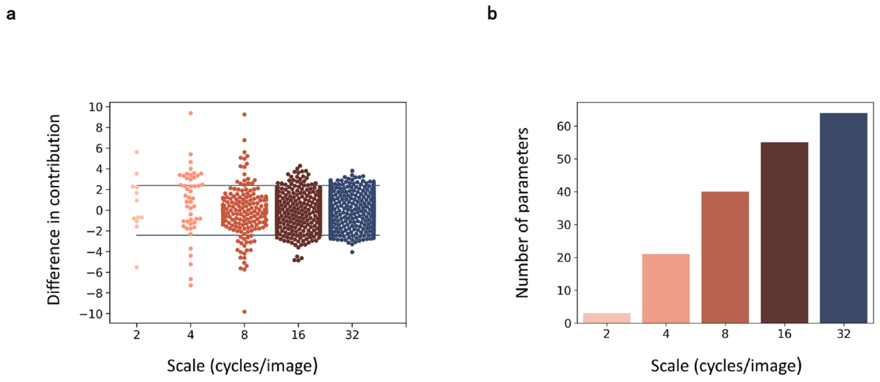


**Figure S2**. (a) Distribution of the contribution difference (the difference between standardized Cohen's d in VGG-Face and humans) at different scales of spatial frequencies. The blue lines denote 1.96 standard deviations from the mean difference; (b) the number of parameters contributing significantly different between the VGG-Face and human observers at different scales of spatial frequencies.

**Supplementary analysis 3**

VGG-16 has the same architecture as the VGG-Face, but different prior task experience as it is pre-trained for object categorization as AlexNet[1]. We trained the VGG-16 to perform the gender classification task with the same transfer learning procedure as that for the VGG-Face, and the testing accuracy of gender classification was 94%, indicating that it was able to perform the task. However, the CI obtained from the VGG-16 (Figure S3) was in sharp contrast to the CIs of human observers either as a whole ($r = 0.14$) or at different scales ($r = 0.44$, -0.05, -0.09, 0.01 and 0.0) at the scales of 2, 4, 8, 16 and 32). We also reconstructed female and male prototypes of VGG-16, and they appeared quite distinct from those of human observers and the VGG-Face (Figure 1c).

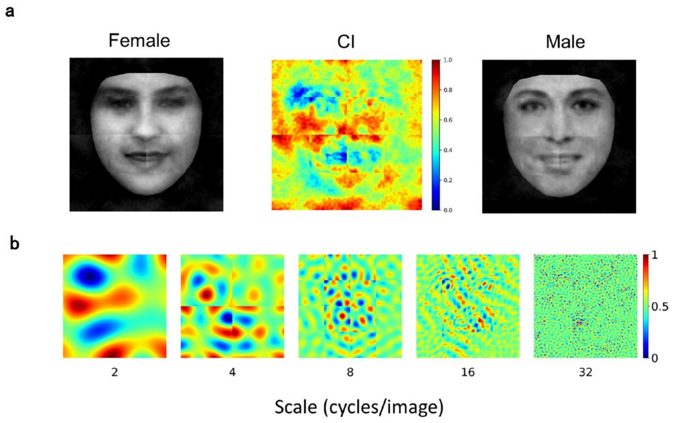


**Figure S3**. Classification images of the VGG-16 as a whole (a) and at different scales of spatial frequencies (b). The faces shown in the figure were synthetic ones created by a morph algorithm and thus had no relation to real human.

**References**

1. Krizhevsky, A., Sutskever, I. & Hinton, G.E. Imagenet classification with deep convolutional neural networks. in *Advances in neural information processing systems* 1097-1105 (2012).
